# Catalytic pocket-informed augmentation of enzyme kinetic parameters prediction via hierarchical graph learning

**DOI:** 10.1101/2025.05.18.654694

**Authors:** Ding Luo, Xiaoyang Qu, Binju Wang

**Author notes:** **Corresponding Author:** (Binju Wang).

## Abstract

Predicting enzyme kinetic parameters is crucial for enzyme engineering and mining, but development of accurate tools for this task is still challenging due to the complexity of influencing factors. Large language models (LLMs)-based deep learning methods show satisfactory performance in Kcat prediction by encoding substrate and enzyme sequence information, but this further enhancement of the performance can be largely limited by the ignorance of the substrate-binding information. Here, we introduce GraphKcat, a deep learning framework that integrates enzyme-substrate 3D binding conformations for precise kinetic parameter prediction. The method employs Chai-1 to generate enzyme-substrate complex conformations, followed by a developed multi-scale hierarchical graph neural network to systematically characterize active site features from all-atom (AA) level to coarse-grained (CG) level, which are subsequently fused with LLMs embedded substrate, sequence, and environmental related information for prediction. To enable efficient multimodal feature integration, we proposed a multi modal cross-attention fusion (MMCAF) module for feature alignment and update. Experimental evaluations demonstrate that GraphKcat can effectively identifying catalysis-critical residues, learning sequence similarity-independent features through its multi-scale graph architecture, and maintaining robust performance under low sequence similarity conditions. Critically, GraphKcat captures conserved structural and physicochemical patterns related to enzymatic activity, enabling accurate identification of high-activity enzyme variants and functional mutants even for low-homology targets. These capabilities highlight its potential as a transformative tool for enzymatic activity prediction, rational enzyme engineering, and enzyme mining in industrial applications.

## Introduction

Enzymes are molecular machines that power nearly all biochemical reactions in living organisms, from digesting food^1^ to replicating DNA^2^. Beyond biology, they are crucial tools in industries such as pharmaceuticals (e.g., synthesizing antibiotics^3^), biofuels (e.g., breaking down plant biomass^4^), and environmental remediation (e.g., degrading plastic waste^5^). However, natural enzymes often lack the efficiency, stability, activity required for industrial applications.^6, 7^ To address this, enzymes engineering (involves designing or modifying enzymes through genetic or protein engineering to enhance their properties)^8, 9, 10^ or enzyme mining (focuses on discovering novel enzymes from natural environments)^11, 12, 13^ strategies were often used to improve the enzyme properties or mining new enzymes with desired properties. In enzyme activity engineering process, determining how enzyme sequence changes affect enzyme behavior, particularly kinetic parameters like catalytic rate (Kcat) and substrate affinity (Km), is critical for accelerating this optimization process.^14, 15^ However, enzyme activity via experimental techniques remains labor-intensive and low-throughput, particularly for large-scale enzyme variant screening.^16, 17^

Recent advances in deep learning have enabled data-driven predictions. Early machine learning approaches in enzyme kinetics prediction predominantly relied on handcrafted features to represent enzymatic reaction processes.^18^ These models employed traditional machine learning algorithms to predict kinetic parameters, or utilizing convolutional networks for modeling one-hot protein sequence representation, or applying graph neural networks to model substrate structures.^19^ However, such simplistic representation paradigms have demonstrated limited generalizability, particularly under test conditions involving unseen data or sequences with low similarity to training datasets.^20^ In contrast, the Transformer architecture, a groundbreaking innovation in natural language processing (NLP), achieves superior semantic representation through its attention mechanisms and mask strategies, enabling the capture of high-order contextual relationships.^21^ This paradigm has been successfully adapted to biological systems, like pretrained Transformer models trained on billions of protein sequences^22^ or small molecules^23, 24^, and exhibit exceptional performance in tasks such as structural prediction and property estimation.^22, 25^ By encoding enzyme sequences and substrate molecules via large language models (LLMs), recent studies have validated enhanced generalization capabilities in predicting enzymatic kinetic parameters compared to conventional feature-engineered models.^26, 27, 28, 29^

While selecting appropriate representation features for enzyme sequences and substrates remains foundational, recent studies suggest that incorporating additional enzyme-reaction-related information or multimodal data might enhance model generalizability. For instance, integrating environmental factors such as temperature, pH, or organism has empirically improved predictive performance.^26, 30^ However, attempts to leverage multimodal language models for joint representation of enzyme sequence-structure relationships have yielded only limited improvement in model performance.^28, 29^ These findings underscore the need for cautious and architecturally refined approaches when fusing heterogeneous, multimodal features, as simplistic integration strategies may fail to capture synergistic interactions. Notably, enzymatic catalysis occurs within spatially defined active sites, recent studies have realized the significant influence of active site information on enzyme catalytic reactions and have attempted to leverage such information to provide prior bias for training enzyme kinetic parameters prediction models.^31^ However, current predictive models for enzymatic activity rarely explicitly incorporate the substrate-binding information, which are highly relevant to both the activity and selectivity of the enzyme catalysis. This gap indicates an underexplored opportunity to enhance predictive models by incorporating the interactions of substrate’s binding to its catalytic pockets.

To address this challenge, we propose a framework called GraphKcat. In addition to the information from enzyme’s and substrate’s and sequences, GraphKcat incorporates the three-dimensional enzyme-substrate active site information. The proposed framework employs a complex conformation prediction tool named Chai-1^32^ to predict enzyme-substrate conformations from enzyme sequences and substrate SMILES. Following active pocket extraction from the predicted conformations, GraphKcat constructs a multi-scale graph neural network to progressively update the encoding of enzyme active pocket information, subsequently integrating these features with LLMs representations of substrates, enzyme sequences, and environmental factors for enzyme kinetic parameters prediction. To enhance the fusion performance of active pocket information and LLMs representations, we design a multimodal cross-attention fusion (MMCAF) module that aligns features across different latent spaces. Demonstrated through comprehensive evaluations, GraphKcat exhibits superior performance attributable to its well-architected model framework and incorporation of critical catalytic pocket characteristics and environmental factors.

## Results

### Structure-based enzyme-substrate representation

GraphKcat encodes complex information of enzyme-substrate pairs. The binding conformations of enzyme complexes were predicted using Chai-1,^32^ a state-of-the-art biomolecular interaction prediction model. Previous studies have demonstrated that Chai-1’s side chain interaction distances align more closely with the reference distribution of native side chains.^33^ To address potential structural clashes in predictions, we employed the FastRelax^34^ module in Rosetta to generate chemically valid structures (**Fig. 1a**). To enhance active site representation, we developed a multi-scale graph neural network (GNN) that models enzyme-substrate reaction sites across resolutions, spanning from the all-atom (AA) level to the coarse-grained (CG) level. In the all-atom graph construction, heavy atoms from both protein and substrate serve as nodes, with edges determined by interatomic distances. For CG-level representation, protein atoms in active sites naturally correspond to amino acid-level nodes. However, substrate atoms exhibit greater structural diversity than amino acids and lack predefined grouping rules. To address this, we adapted subgraph extraction strategies from PS-VAE,^35^ systematically dividing substrate atoms into CG nodes with explicitly defined edges (**Fig. 1b**). Each CG node in the substrate representation is a molecular fragment containing multiple atoms.

**Fig. 1:**
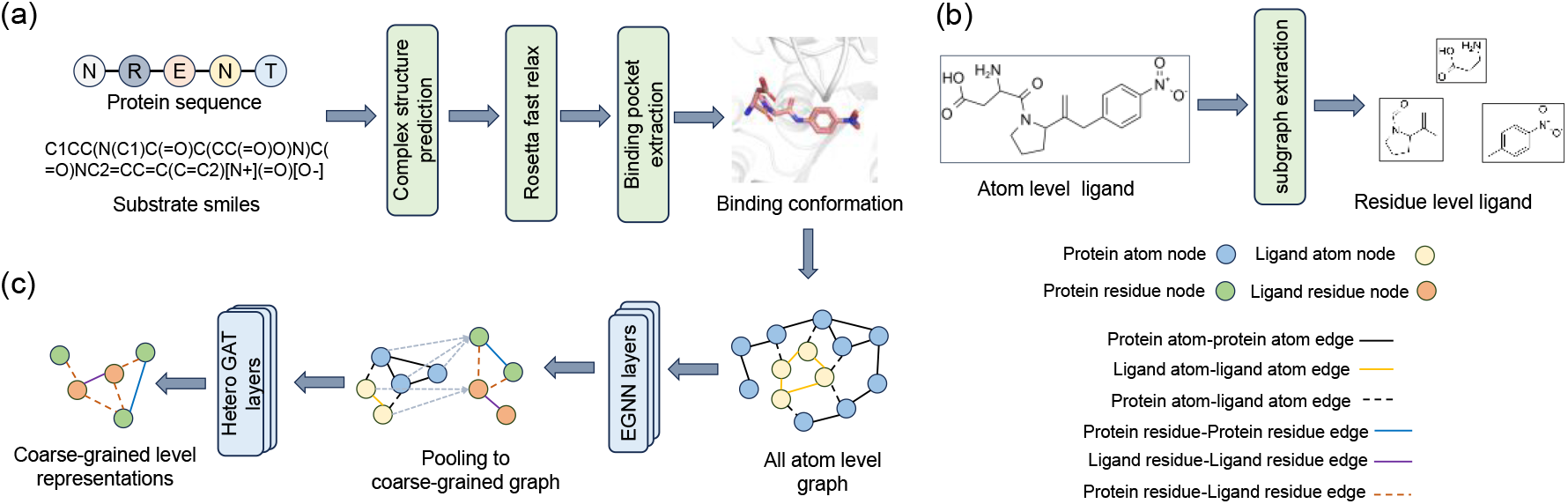
Graph encoder architecture for enzyme-substrate active site representation. (a) Protein sequences and substrate SMILES are processed through Chai-1 for complex structure prediction, followed by energy minimization using Rosetta’s FastRelax module. Binding sites are identified through substrate spatial positioning analysis. (b) Substrate atoms are systematically clustered into coarse-grained (CG) nodes using PS-VAE-derived fragmentation criteria. (c) The binding site is converted into an all-atom graph processed through E(3)-equivariant GNN layers, with atomic features being aggregated into CG graphs for subsequent heterogeneous GATv2Conv-based updates (see **Methods** section).

The dual-scale graph captures distinct modal information of enzymatic reaction active sites. We implemented SE(3)-equivariant graph neural networks (EGNN)^36^ for updating node features in the AA graph representation (**Fig. 1c**). These AA graph embeddings encode atomic-level interaction features between enzyme and substrate. Through atom indexing that maps individual atoms to corresponding coarse-grained (CG) nodes, these atomic features were aggregated via a single directed GATv2Conv layer.^37^ The resulting pooled CG features were subsequently combined with initial CG features to construct the complete CG graph. Given the distinct nature of pocket CG nodes and substrate CG nodes, we applied three specialized GATv2Conv layers to respectively handle: 1) intra-molecular interactions among substrate CG nodes, 2) intra-molecular interactions within pocket CG nodes, and 3) inter-molecular interactions between substrate and pocket CG nodes (see **Methods** section). It is worth noting that recent studies have demonstrated the dependency of complex-based graph encoders’ generalization performance on their architectural design.^38, 39^ To validate the superiority of our developed GNN architecture, we trained a protein-ligand binding affinity prediction model using this framework on PDBbind dataset^40^ and benchmarked it against existing methods on the CASF-16,^41^ which are the most widely used standard training set and golden test set in this task. Our GNN architecture demonstrated comparable performance with other state-of-the-art (SoTA) models (**Supplementary Table S1**), confirming its robust capability for active site information extraction. This hierarchical multi-scale architecture enables efficient and rational aggregation of multimodal active site information, significantly improving enzymatic reaction representation for kinetic parameter prediction.

### Model overview of GraphKcat

GraphKcat is a multi-modal deep learning framework designed to predict enzyme kinetic parameters Kcat and Km through integrated sequence-structure information (**Fig. 2a**). Leveraging large language models (LLMs) for sequence and substrate representation, we capitalize on their unsupervised pre-training across billion-scale datasets to capture high-order patterns in substrate and enzyme sequences. The substrate’s atomic-level representation derives from Uni-Mol2^24^, a multi-modal foundation model pre-trained on large molecular datasets. Crucially, graph encoder outputs and LLM embeddings require alignment within a unified latent space to synergize their complementary strengths. Our multi-modal cross-attention fusion (MMCAF) architecture addresses this integration challenge, with fused embeddings further combined with organism-specific features, reaction temperature, and pH values. Finally, two linear layers independently predict logKm and logKcat values.

**Fig. 2:**
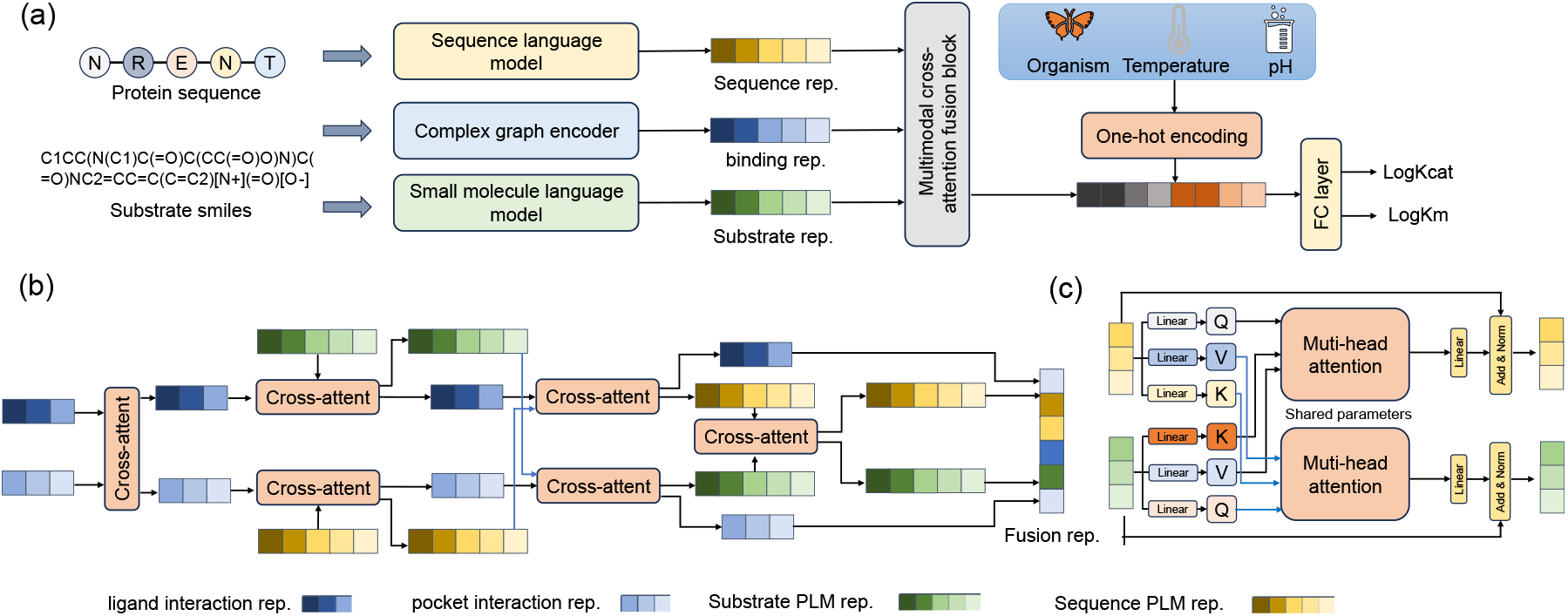
The overview model architecture of GraphKcat. (a) The model predicts Kcat/Km values through synergistic integration of four data modalities: substrate language embeddings, protein sequence embeddings, enzyme-substrate complex graph features, and environmental parameters (organism, temperature, pH). Our multi-modal cross-attention fusion module integrates language representations with structural features through learned attention patterns. (b) The fusion module employs six parameter-shared cross-attention units that iteratively refine feature interactions between modalities, while residual connections preserve original embedding information throughout the fusion process. (c) Each cross-attention unit implements multi-head attention mechanisms to mutually update input features, establishing bidirectional information flow between active site structural and sequences representations (see **Methods** section).

The MMCAF block mainly consist of six cross-attention modules. Those cross-attention blocks enable to fusion the active site embeddings with LLM embeddings (**Fig. 2b**). According to the CG node index, we split the CG graph nodes to pocket nodes and substrate nodes. Those nodes were interacted each other or interact with LLM residue level embeddings by cross-attention module. The cross-attention module calculates the attention weights between query embeddings and key embeddings to weight sum the feature’s importance for the final embeddings (See **Methods** section). This hierarchical attention framework establishes context-aware modality integration, significantly enhancing enzymatic reaction modeling through learned complementarity between structural and LLMs sequential descriptors.

### Active site information improves the generalization of Kcat/Km prediction

Our comparative analysis evaluated the impact of protein language model selection on prediction performance, comparing sequence-based ESM2_3B^22^ against structure-aware SaProt (SaProt_AF2_650M),^42^ both integrated with Uni-Mol2 v1.1B’s molecular embeddings (**Fig. 3**). Across all Kcat/Km prediction tasks without sequence similarity filtering, ESM2 demonstrated superior performance (PCC: 0.823 and 0.780 Km; RMSE: 0.855/0.941) over SaProt, suggesting limited predictive value from structural information in general enzyme modeling scenarios, which is consistent with prior studies.^28^ However, under strict sequence dissimilarity conditions (<40% identity between training/test sets), SaProt exhibited marginally better generalization for Kcat prediction (PCC: 0.412 vs 0.389; RMSE: 1.603 vs 1.597), while showing comparable Km prediction accuracy (PCC: 0.667 vs 0.667; RMSE: 0.946 vs 0.927). Those results imply that using model with more parameters (ESM2_3B) and extract modality information (SaProt_AF2_650M) are both important for improve model performance. The performance with or without considering similarity cutoff between training and test set are both crucial, as they reflect the enzyme mutation prediction and enzyme miner task in real scenarios. Given balanced consideration of both operational paradigms, ESM2 was selected as the primary sequence embedding provider due to its robust overall performance.

**Fig. 3.**
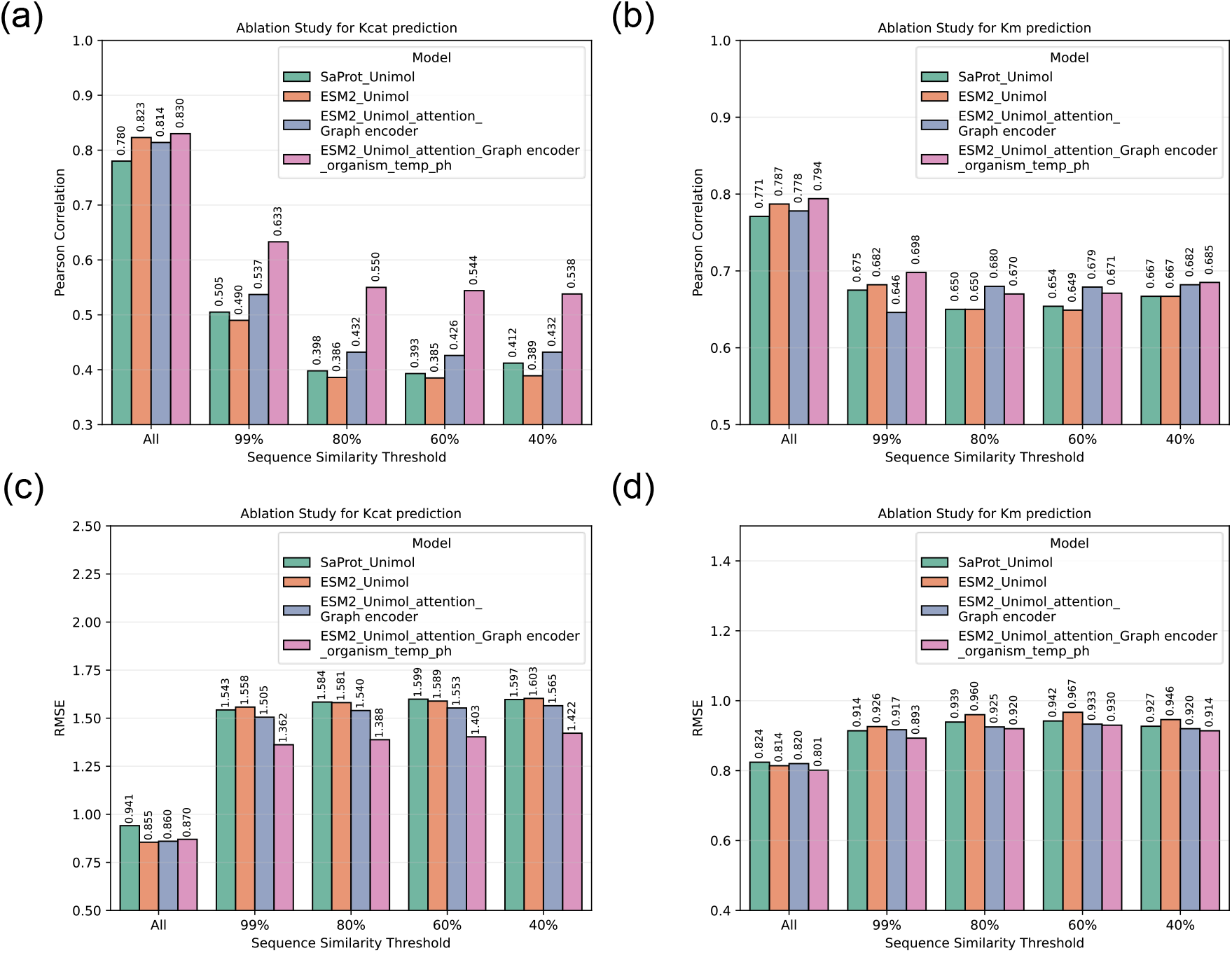
Ablation analysis of GraphKcat performance under diminishing enzyme sequence similarity to the training set. (a-b) Pearson correlation coefficient (PCC) between predicted and experimental kcat values across sequence similarity intervals (100-40%, left to right) for (a) Kcat (b) Km (c-d) Corresponding root mean square error (RMSE) progression. Model variants include: SaProt_Unimol: Baseline architecture using SaProt structural features and Unimol molecular representations; ESM2_Unimol: Enhanced version incorporating ESM2 evolutionary-scale embeddings with Unimol features; ESM2_Unimol_attention_Graph encoder (square markers): Extended framework integrating ESM2 embeddings, Unimol molecular graphs, and active site information processed through multimodal cross-attention fusion; ESM2_Unimol_attention_Graph encoder_organism_temp_ph: Full implementation with additional organism-specific environmental predictors (temperature/pH adaptation).

Our ablation studies (**Fig. 3**) reveal differential impacts of structural components across experimental conditions. Under standard evaluation protocols (no sequence similarity constraints), integration of active site features through the MMCAF framework shows comparable Kcat prediction performance to language-only baselines UniKP (ΔPCC <0.02). However, under strict sequence dissimilarity (≤40% identity), architectural integration of graph-encoded active site information yields significant generalization improvements: Kcat prediction achieves PCC=0.432 (RMSE=1.565) and Km prediction reaches PCC=0.682 (RMSE=0.920), surpassing language-only models by 5.6% and 2.2% in PCC respectively. Notably, Km prediction demonstrates greater performance gains from active site features, aligning with biochemical theory that substrate binding affinity (Km’s determinant) depends critically on active site geometry, whereas Kcat involves dynamics including transition-state stabilization and product release kinetics. This performance underscores two fundamental insights: 1) structural features of active pocket provide complementary predictive power under evolutionarily distant enzyme scenarios, partially compensating for sequence representation limitations; 2) Pure LLMs embeddings struggle to capture the intrinsic characteristics about enzyme reactions, as pure LLMs embedding may implicitly learn limited enzyme reaction information in sparse datasets saturation. These findings validate our multi-modal approach’s superiority in modeling real-world enzyme engineering challenges.

Notably, the incorporation of environmental related features including organism, pH, and temperature further enhances model performance in both Kcat and Km predictions. Quantitative analysis reveals consistent improvements in Kcat prediction accuracy across varying sequence similarity thresholds (100-40%), suggesting these features provide extra information to sequence data that independently contributes to enzymatic turnover rates. In contrast, the predictive gains for Km parameters decrease progressively with decreasing sequence similarity between training and test datasets. This differential performance aligns with the distinct mechanistic determining factors of each parameter, as Km predominantly reflects substrate binding affinity within the enzyme’s active site, demonstrating weak correlation with unrelated biochemical factors, Kcat appears more sensitive to broader environmental and organismal context.

### Compare GraphKcat with other deep learning models

Our benchmark evaluation compares GraphKcat against established deep learning approaches in enzymatic parameters prediction (**Fig. 4a-b** for PCC values, **Supplementary Fig. 1** for RMSE values). DLKcat,^19^ the first deep learning framework for Kcat estimation, employs graph neural networks (GNNs) for substrate modeling and convolutional neural networks (CNNs) for enzyme sequence processing. Comparative baselines also include UniKP^26^ and MPEK,^30^ which are LLMs driven approaches where UniKP combines Extra Trees regression with molecular embeddings, while MPEK enhances predictions through environmental feature integration (organism source, pH, temperature) via multilayer perceptions (MLPs). All models underwent rigorous training and evaluation on identical benchmark datasets under standardized conditions.

**Fig. 4.**
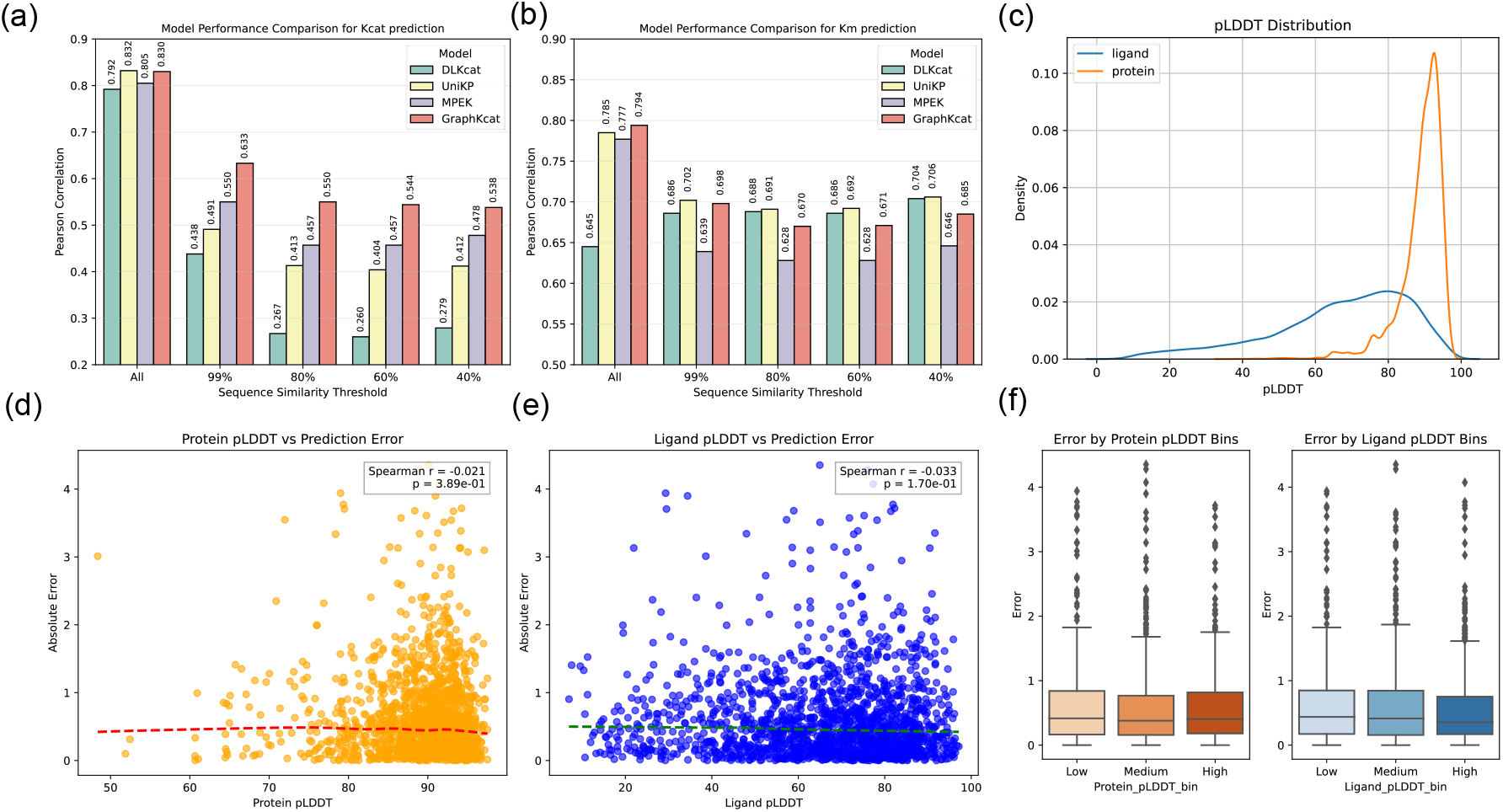
Comparative model performance and structural quality impacts of predictive efficacy. (a-b) Pearson correlation coefficients (PCC) for Kcat (a) and Km (b) predictions across DLKcat, UniKP, MPEK, and GraphKcat models under declining enzyme sequence similarity thresholds (100-40%) relative to the training set. All models were trained and evaluated on identical datasets. (c) Structural confidence metrics (pLDDT distributions) for enzyme-substrate complexes predicted by chai-1. (d-e) Spearman correlations between pLDDT values (d) and absolute prediction errors for protein (e) and ligand for Kcat values prediction in test datasets. (f) Prediction error distributions stratified by low/medium/high protein-ligand pLDDT confidence.

The results demonstrated that GraphKcat significantly outperformed other methods, particularly when sequence similarity between training and test sets decreased. Without considering sequence similarity, DLkcat, UniKP, MPEK, and GraphKcat achieved PCC values of 0.792, 0.832, 0.805, and 0.830 respectively for Kcat prediction, and 0.645, 0.785, 0.777, and 0.794 for Km prediction. DLkcat exhibited slightly worse performance compared to other methods, likely because alternative models all utilized language model embeddings as input features, which have been proven advantageous in diverse biomolecule-related tasks.^22, 25^ UniKP employed classical machine learning algorithms and demonstrated comparable performance to GraphKcat in both Kcat and Km predictions. This aligns with the understanding that traditional machine learning methods possess stronger data-fitting capabilities than deep learning approaches when working with small datasets. However, under more rigorous evaluation conditions where sequence similarity between training and test sets was reduced to 40%, GraphKcat maintained competitive performance with PCC values of 0.538 (Kcat) and 0.685 (Km). These results surpassed those of UniKP’s Kcat prediction (0.412) and DLkcat’s Kcat prediction (0.279), and outperformed MPEK (0.478 for Kcat, 0.646 for Km). This observation proves previous findings that DLkcat struggles to predict meaningful Kcat values for mutant and unfamiliar enzymes.^20^ Notably, DLkcat demonstrated better generalizability for Km prediction under reduced sequence similarity (40% similarity: 0.704 PCC), though its performance remained limited without sequence similarity constraints (0.645 PCC). This contrast between DLkcat’s Km and Kcat prediction capabilities under varying conditions worths further investigation. Overall, GraphKcat exhibited superior robustness compared to other deep learning methods across both Kcat and Km prediction tasks.

### GraphKcat is not sensitive to the quality of input structures

Compared to conventional Kcat/Km prediction models, GraphKcat uniquely requires enzyme-substrate complex structural inputs. While such complexes can be computationally generated via docking or co-folding algorithms (e.g., AlphaFold3^43^), achieving high-confidence structural predictions remains not easily in some cases.^44^ We systematically evaluated structural confidence of training and test set using predicted Local Distance Difference Test (pLDDT) metrics (**Fig. 4c**). Enzyme structures exhibited consistently high pLDDT scores, indicating reliable protein conformation predictions, while substrate pLDDT values remained within acceptable thresholds suggesting moderate but functionally sufficient ligand pose accuracy.

To quantify structural confidence impacts on predictive performance, we conducted Spearman correlation (ρ) analyses between pLDDT scores and absolute prediction errors. As shown in **Figs. 4d-e, Supplementary S3**, neither protein (ρ = -0.021, p=3.89e-01) nor substrate (ρ = -0.033, p=1.70e-01) pLDDT demonstrated significant linear correlations with prediction errors in Kcat prediction, the Km values prediction task shows the same results (**Supplementary. Fig. S2-S3**). Stratification of test cases into low/medium/high pLDDT bins further demonstrated that prediction errors remained statistically indistinguishable across different confidence levels. While substrates in high-pLDDT bins exhibited a marginal reduction in mean absolute error, but trend failed to reach statistical significance (**Fig. 4f, Supplementary Fig.S3**). These results demonstrate that GraphKcat maintains prediction robustness across broad structural quality variations. The model’s insensitivity to input structural accuracy suggests inherent capacity to extract functionally relevant features even from imperfect active site binding conformations, which is a critical advantage for applications where experimental structures is unavailable. This structural tolerance, coupled with competitive performance metrics (**Figs. 4a-b**), proves GraphKcat as a generalizable solution for enzymatic parameter prediction.

### GraphKcat can distinguish high activity mutations and sequences

We next evaluated GraphKcat’s performance in enzyme mining and mutational effect prediction tasks. Tyrosine ammonia-lyase (TAL),^45^ a critical biocatalyst for synthesizing aromatic compounds including flavonoids, cinnamoyl anthranilates, requires continuous discovery of novel variants and activity optimization through protein engineering. We benchmarked GraphKcat using two literature-curated TAL datasets: 1) a homolog mining set containing the reference IsTAL sequence with relatively low native activity compared to other homologs, and 2) a mutational screening set comprising engineered variants (MT-587V, MT-10Y, MT-489T with enhanced catalytic efficiency, and five efficiency-reducing mutants), those datasets were also be utilized to benchmark other kinetic parameters prediction tools (UniKP, MPEK, CatPro), The predicted structural confidence can be found in **Supplementary Table S2**. As shown in **Fig. 5a-b**, prediction success was determined by correct identification of activity trends relative to reference samples. Green markers denote accurate predictions of increased/decreased Kcat/Km ratios, while orange indicates wrong prediction results. GraphKcat demonstrated 86% accuracy (5/6 high-activity homologs correctly identified) in mining tasks and 55% overall accuracy in mutation screening, with successful trend prediction for 3/4 efficiency-enhancing mutants. These results highlight GraphKcat’s potential for identifying catalytically superior enzyme variants through both natural sequence mining and rational mutagenesis approaches.

**Fig. 5.**
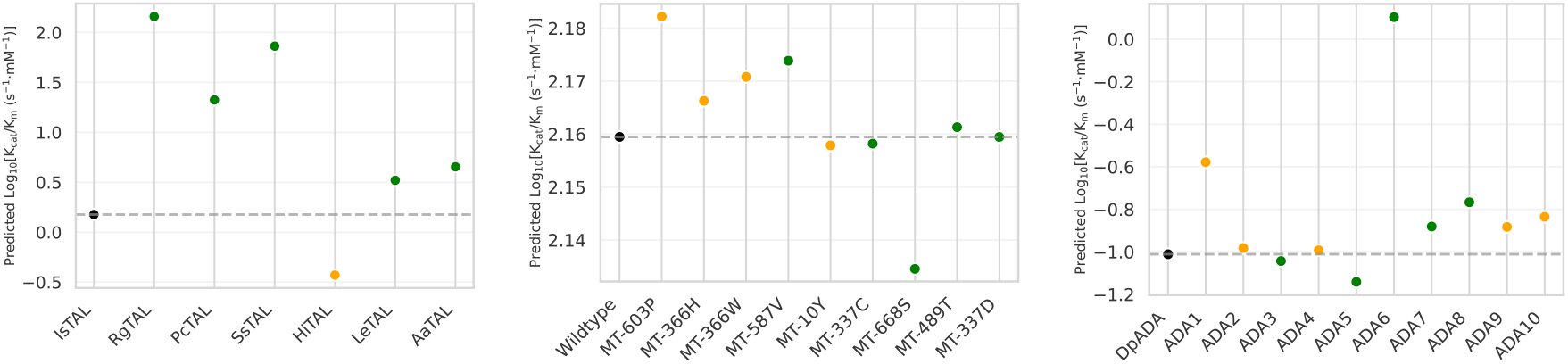
Performance evaluation of GraphKcat in enzyme mining and mutation prediction tasks. (a) Predicted versus experimental Kcat/Km values for transaminase (TAL) enzyme mining, (b) TAL mutation screening, and (c) D-amino acid deaminase (DpADA) mining tasks. Black dots indicate reference values, while green and orange dots represent correct and incorrect predictions, respectively (correctness determined by whether the model accurately identified activity increases/decreases relative to reference samples).

Subsequent evaluation extended to acetaldehyde dehydrogenase (DpADA) mining tasks, where GraphKcat demonstrated robust generalization capabilities. DpADA enzymes catalyze the critical conversion of toxic acetaldehyde, a volatile organic compound (VOC) to acetyl-CoA, a versatile metabolic precursor with broad industrial applications.^46^ This transformation represents an essential step in developing sustainable bioprocesses for VOC valorization. When benchmarked against the ADA homolog dataset (ADA1-10) curated by Zhou et al.,^47^ which identified ADA6, ADA7, and ADA8 as high-activity variants (3 to 7-fold catalytic improvement over ADA), GraphKcat successfully recognized all three superior candidates (**Fig. 5c**). Notably, these predictions were achieved despite low sequence homology between the test sequences and training set enzymes (all pairwise identities ≤60%), underscoring the model’s capacity to extract functional patterns beyond primary sequence conservation. The agreement between GraphKcat’s predictions and experimental tests demonstrates its ability to predict enzyme activity accurately even when dealing with evolutionarily distant enzymes.

### Attention heatmaps highlight catalytic hotspots

To validate whether attention patterns reflect biologically meaningful features, we analyzed how GraphKcat links enzyme active sites to their substrates. We investigated two key digestive enzymes using GraphKcat’s attention maps to decipher their functional patterns: carboxylesterase (EC 3.1.1.1), a serine hydrolase that hydrolyzes ester bonds in neurotransmitters, and β-galactosidase (EC 3.2.1.23), a glycosidase essential for lactose metabolism. Spatial attention heatmaps showed strong correlations with established catalytic mechanisms. For carboxylesterase, elevated attention weights (scores > 0.05) precisely aligned with the catalytic triad (Ser144-His295-Asp266), while Tyr72—a residue mediating hydrophobic interactions with the substrate was also identified. In β-galactosidase, three conserved glutamate residues (Glu201/Glu417/Glu504) were mapped to the active site, as confirmed by multi-sequence alignment (MSA) analysis (**Supplementary Fig. S5**). Attention peaks distinctly highlighted the nucleophilic Glu201, a pivotal residue for the double-displacement mechanism, along with galactose-binding subsites (Arg109/Asn200/Asn421/Glu474) that stabilize the substrate’s chair conformation. These results demonstrate that GraphKcat’s attention weights inherently capture both catalytic residues and functionally critical binding architectures across diverse enzyme classes.

## Discussions

The enzyme kinetic parameters Kcat and Km are essential metrics for quantifying enzymatic activity and elucidating catalytic mechanisms. It provides critical insights into the rate-limiting step of the reaction and serves as a benchmark for comparing enzyme performance. These parameters are fundamental in rational enzyme engineering, drug discovery, and metabolic flux analysis. Although several deep learning strategies has been utilized to predict the enzyme kinetic parameters, but those models were not trained with explicit enzyme reaction active site information, that information are curial for enzyme kinetic parameters, thus lead to limited generalization ability. GraphKcat considered the detailed enzyme reaction active site information from all-atom level to coarse-grained level by a carefully designed multi-scale geometric deep learning framework (**Fig. 1**) for the first time, and showed completive performance with other deep learning methods in same training and test set (**Fig.4a-b**).

Enzyme kinetic parameters are influenced by complex factors. To train a robust deep learning model for predicting these parameters, it’s important to consider as many relevant parameters as possible and to reasonably integrate information from different modalities. The performance of these models in downstream tasks depends directly on how well the representations from different modalities are aligned. In GraphKcat, we developed an MMCAF block to align the large language model embeddings with geometric graph embeddings (**Fig. 2b**). Ablation studies show that this fusion strategy improves the model’s ability to generalize, even when the sequence similarity between the training and test sets drops to 40% (**Fig. 3**). This finding is also evident in the attention map of the MMCAF module (**Fig. 6**). The model captures information about enzyme catalysis in the active pocket, suggesting it may be learning aspects of enzyme catalysis beyond just sequence similarity. We also discovered that environmental factors play a key role in predicting Kcat and Km, as these factors are independent of sequence similarity and affect enzyme parameters. Additionally, we assessed the impact of the enzyme-substrate complex quality on prediction error, since high-quality structures are not always available. Our results revealed no significant correlation between the quality of enzyme or substrate structures and the prediction error. This demonstrates that GraphKcat’s performance remains robust regardless of the quality of input structures, making it a valuable tool for enzyme-related tasks.

**Fig. 6.**
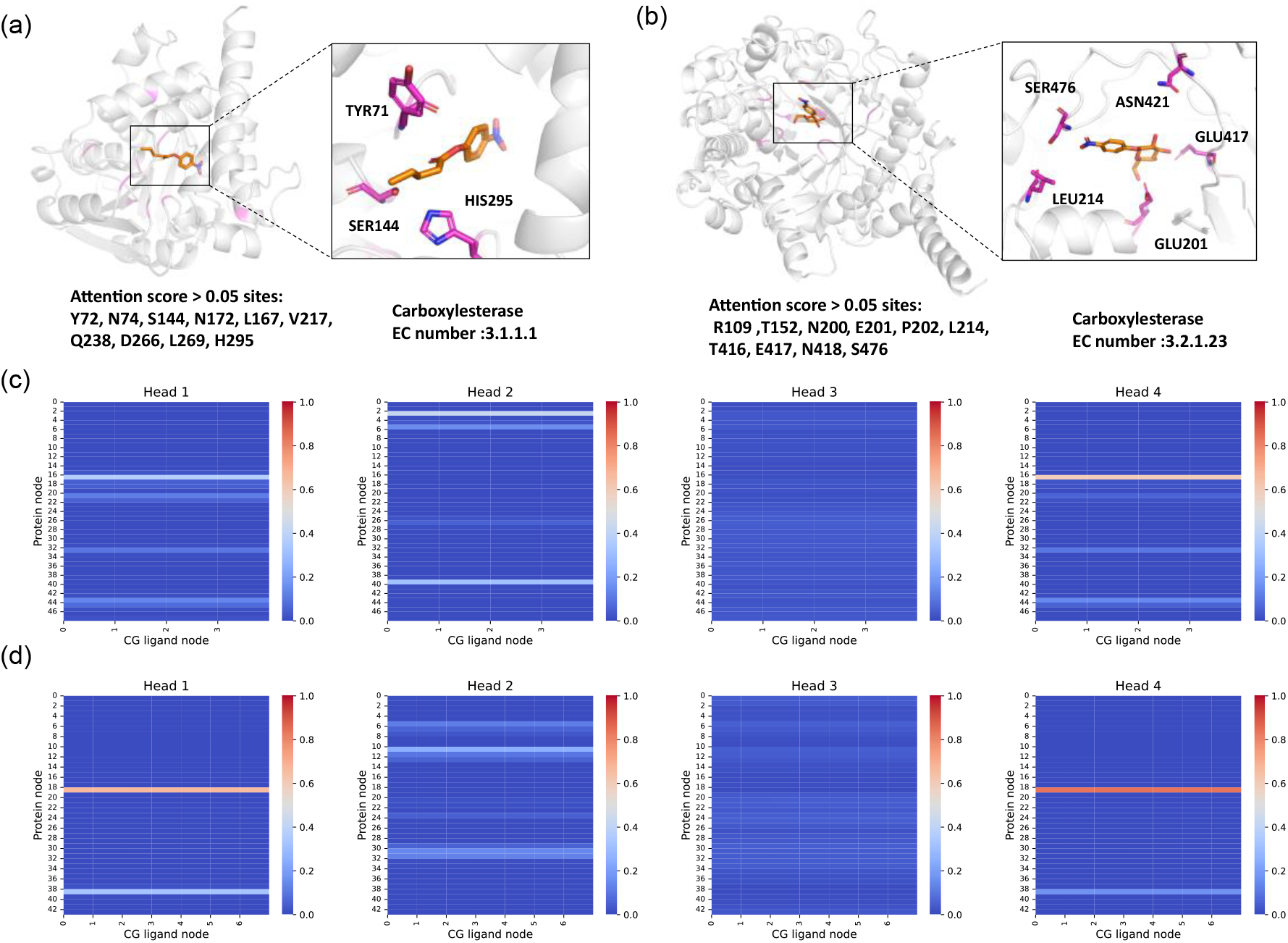
Attention analysis between enzyme active pockets and ligands. (a, b) Substrate-binding structures of carboxylesterase (a) and β-galactosidase (b). Magenta highlights amino acids in the active pocket with attention scores >0.05. Right insets zoom into key amino acids (within 4 Å of the ligand) that show high attention scores (>0.05). (c, d) Attention maps between the active pocket (rows) and ligand atoms (columns) for carboxylesterase (c) and β-galactosidase (d). Darker red indicates stronger attention weights.

Enzyme mining tasks and mutation effect prediction in the TAL enzyme dataset and DpADA enzyme dataset demonstrate that GraphKcat can identify highly active sequences and mutations. Compared to reference sequences, GraphKcat successfully predicted five of the six highly active sequences in the TAL enzyme dataset and all highly active sequences in the ADA enzyme dataset. For mutation effect prediction, GraphKcat accurately identified three of the four highly active mutants. However, its performance in predicting low-activity sequences still requires improvement. This limitation may arise because all training data contain sequences with measurable enzyme parameters, while the model has not been exposed to sequences or mutants exhibiting negligible or extremely low activity. This saturation effect is also observed in structure-based protein-ligand binding affinity prediction tasks.^48^ Enhancing the training set with negative samples of low-activity or inactive enzyme sequences and mutants could potentially refine the model’s ability to distinguish activity levels, thereby improving prediction accuracy across the full activity range.^49^

We have demonstrated that rational incorporation of enzyme catalytic pocket information can significantly enhance the performance of deep learning models in predicting enzyme kinetic parameters. However, the information currently considered by existing models remains insufficient to fully characterize the complexity of enzyme catalytic reaction processes. Most enzymes require metals or cofactors to generate active intermediates for substrate catalysis during reactions, and such information plays a crucial role in determining enzymatic activity. Unfortunately, current deep learning models have not incorporated these factors due to the absence of comprehensive datasets containing complete reaction information. While GraphKcat integrates structural features of enzyme active pockets, it faces inherent limitations because enzyme catalysis is a highly dynamic process involving substrate entry, binding, intermediate transformation, and product release. A single static structural representation cannot fully capture the transient states and conformational changes, resulting in constrained predictive performance. Evolutionary information encoding offers potential solutions for modeling dynamic aspects of protein function.^50^ Although LLMs implicitly learn evolutionary patterns from natural sequence variation, explicit modeling of evolutionary constraints through methods like MSA has demonstrated superior performance in downstream prediction and generation tasks.^43, 51^ Notably, MSA-guided approaches have successfully engineered enzymes with natural catalytic functions, providing direct evidence that evolutionary information encodes features related to enzymatic activity.^52, 53^ Therefore, systematic integration of evolutionary information might enhance model generalization by bridging the gap between static structural representations and dynamic catalytic mechanisms.

## Methods

### Dataset construction

The MPEK^30^ training and test sets were used for model development, containing 17,893 kcat entries and 24,585 Km entries respectively. To incorporate active site information for each enzyme-substrate pair, we employed Chai-1 v0.5.1,^32^ a state-of-the-art protein-ligand co-folding tool, to generate binding poses and protein structures. These structures subsequently underwent Rosetta FastRelax^34^ relaxation to eliminate potential atomic clashes. Entries with failed structure predictions or poses distant from the protein binding pocket were excluded. The final curated dataset comprised 21,999 training entries (13,968 Kcat, 19,129 Km) and 2,774 test entries (1,762 Kcat, 2,414 Km).

Sequence similarity thresholds were established using MMseqs2^54^ to cluster training and test sequences at 40%, 60%, 80%, and 99% identity levels. Filtered test sequences below each similarity threshold were used to benchmark model generalization capabilities.

### Language model embeddings of substrates and proteins

To better represent substrate information, we used Uni-Mol2^24^ (1.1B) to extract atom-level substrate representations. Uni-Mol2 is a multimodal small molecule pretraining model that employs a two-track transformer architecture to integrate features at the atomic level, molecular graph level, and geometric structure level. For protein residue-level representations, two masked protein language models were compared. These included the multimodal language model SaProt^42^ (SaProt_650M_AF2) with 1024-dimensional features, and the masked protein language model ESM2 (esm2_t36_3B_UR50D) with 2560-dimensional features.

### Active site information embeddings

The active site of enzyme reactions was extracted based on predicted substrate positions using an 8 Å pocket-cutoff. To better represent active site information, we proposed a multi-scale graph neural network architecture. An all-atom-level graph (AALG) and a coarse-grained-level graph (CGLG) were constructed. AALG embeddings were transferred to the CGLG through a graph pooling layer to unify multimodal information. Specifically, Specifically, the active site information was initially represented as an AALG *G* = (*v*_aa_, *e*_aa_), where *v*_aa_ represent atom level nodes and *e*_aa_ denotes edges between nodes. Node features consist of one-hot encodings of the atom types (C, N, O, S, F, P, Cl, Br, I, Unknown), atom degree, implicit valence, and hybridization state. Atoms within 5 Å of another atom were connected by edges. We used one-hot encoding of edge types (substrate-substrate, protein-substrate, protein-protein) and atomic distances as edge features **e**_*ij*_. All the detail features can be found in **Supplementary Table S3**. Equivariant Graph Neural Networks (EGNN) were employed to update node features ***h*** and coordinate ***x*** for each atom, using equation 1-3:

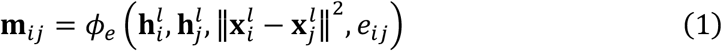

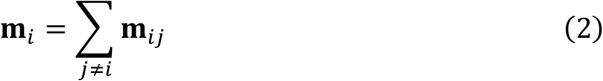

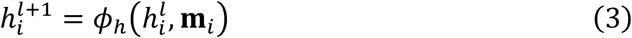

The term **m**_*ij*_ represents the messages between atom ***i*** and atom ***j***. These messages were generated by combining the node features **h**_*i*_, **h**_*j*_, the squared relative distance between atomic coordinates **x**_*i*_, **x**_*j*_, and edge feature **e**_*ij*_ through the edge operation *ϕ*_*e*_ defined in Equation 1. For each node ***i***, the aggregated message **m**_*i*_ was computed by summing messages from all neighboring nodes using Equation 2. Finally, the updated node features 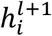 were obtained by processing the aggregated message **m**_*i*_ and the original node feature **h**_*i*_ through the node operation *ϕ*_*h*_.

The CGLG contains residue-level substrate nodes and protein nodes. To create residue-level substrate representations, the substrate was divided into multiple sub-fragments using the subgraph division method from PS-VAE, with a vocabulary size of 350 predefined by the ZINC250K dataset.^55^ For 3D graph construction, the coordinate centroid of all atoms within each subgraph was assigned as the positional embedding for residue-level substrate nodes. The Cα coordinates of protein residues served as residue-level protein nodes. Edges in the CGLG were established between residues located within 8 Å of each other.

We defined two single directed graphs 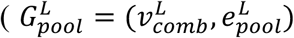, and 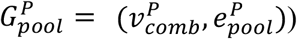 to pool the AALG features into CGLG features using GAT2Conv layer. First, atom-level nodes and predefined residue-level nodes were combined through Equations 4 and 5. Where 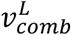 represents the node set comprising substrate atom-level nodes 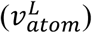 and residue level substrate nodes 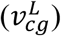. While 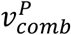 denotes the node set containing protein atom-level nodes 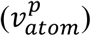 and residue-level protein nodes 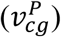. Single directed edges were established from 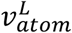 to 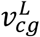 and from 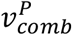 to 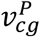 via equation 6 and 7. Batch graphs were implemented to optimize the pooling process, where 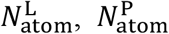 indicate the total number of substrate atoms and protein atoms, respectively. The batch indices Batch_*L*_(*i*), Batch_*p*_(*i*) determines coarse-grained node assignments for substrate and protein atoms.

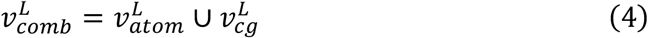

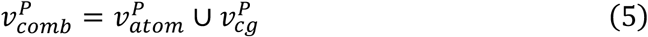

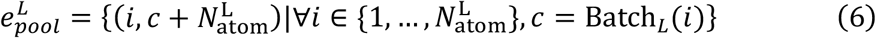

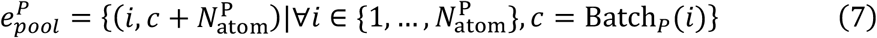

In the aggregation process of GATv2, messages propagated exclusively along the directions defined by edges 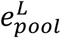 and 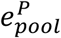. For instance, the attention weights for the substrate atom pooling were calculated via Equation 8, where **a**^⊤^ denotes the attention parameters and, **W** corresponds to learnable weights. Here, 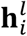 represents the EGNN-updated substrate atom features, while 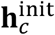 indicates the initial features of the CG node, initialized to zero vectors. Messages from neighboring nodes (𝒩(*c*) = {*i* ∣ (*i, c*) ∈ *e*}) were aggregated to the CG node using the computed attention weights as specified in Equation 9.

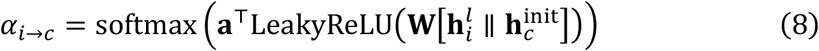

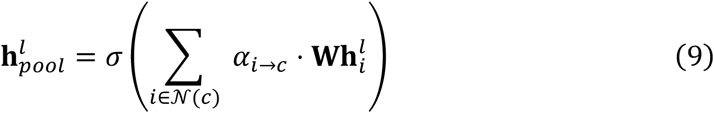

In addition to features pooled from the atom-level graph, he one-hot embedding of the substrate vocabulary size was concatenated with 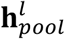 to initialize the substrate coarse-grained (CG) node features. Similarly, one-hot embeddings of amino acid types and backbone dihedral features were concatenated with 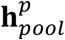 to initialize the protein CG node features. The edge features of CGLG were computed by transforming pairwise distances into radial basis function (RBF) embeddings. This process began by defining a distance range (*d*_*min*_, *d*_*max*_) based on the interaction type—2-22 Å for protein-protein and protein-substrate nodes, and 2-5 Å between substrate CG nodes. Equally spaced centers *d*_*i*_ were generated across these intervals using Equation 10. The width parameter *σ* was derived from Equation 11 to ensure uniform coverage. Each RBF feature dimension *ϕ*_*i*_(*d*) was computed via a Gaussian kernel defined in Equation 12, where *d* represents the input distance. This transformation converted scalar distances into 16-dimensional feature vectors encoding multi-scale spatial relationships through overlapping Gaussian distributions. Protein-protein edge features further incorporated positional embedding features.

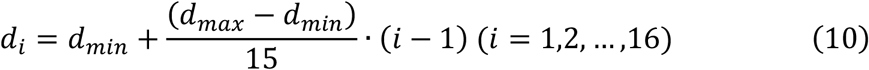

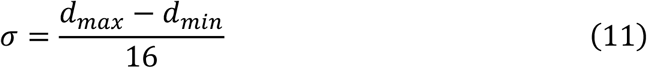

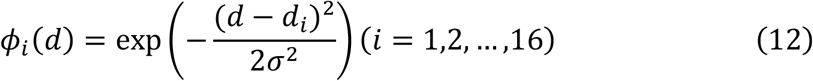

Given the distinct node types between coarse-grained (CG) substrate nodes and CG protein nodes, we implemented heterogeneous message passing layers to update CGLG features. The intra-message passing for substrate CG node features 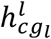 and protein CG nodes 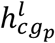 were updated by the GATConv2 layers:

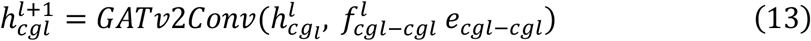

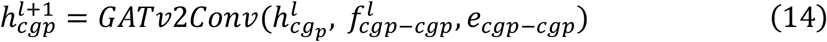

Where 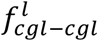 represent the edge features between substrate CG nodes, while 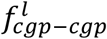 denotes the edge features between protein CG nodes. Substrate CG nodes and protein CG nodes were then updated through defined inter-edges connecting substrate CG nodes to protein CG nodes:

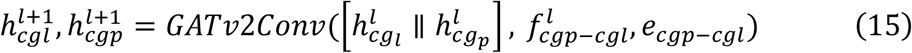

Where 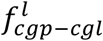 represent the edge features between the substrate CG nodes and protein CG nodes.

### Muti-modal feature fusion block

To better utilize both language model features and active site information, we implemented a multimodal feature fusion block to align these distinct modality features within a shared embedding space. This integration was achieved through cross-attention mechanisms, such as those defined in Equations 16 and 17, which employed parameter-sharing strategies to enable interaction between substrate coarse-grained (CG) features and pocket CG features.

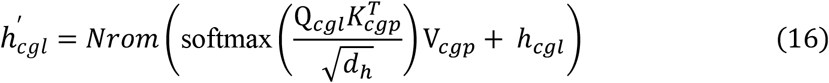

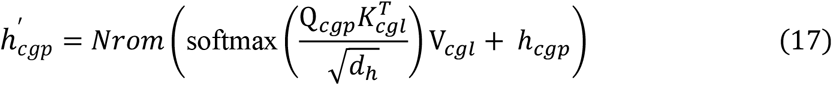

The terms **Q, K, V** denote linear transformations of input features, where *d*_*h*_ represent dimension. Six cross-attention blocks were employed to align the ligand coarse-grained (CG) representations, pocket CG representations, substrate language model embeddings (*h*_*lml*_), and protein sequence language model embeddings represent the residue level language model embedding of substrate (*h*_*lmp*_) through Equations 18-23. The final updated LLM representations, pocket CG representations, and ligand CG representations, were concatenated and processed by two dedicated linear layers for Kcat and Km prediction. Both tasks employed MSE loss with equal loss weights of 0.5 each.

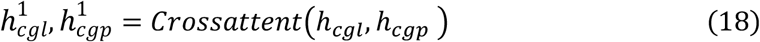

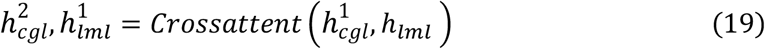

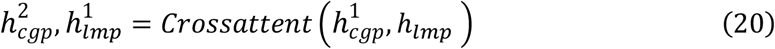

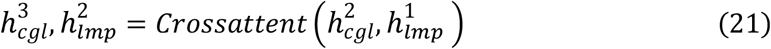

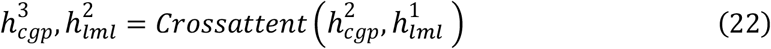

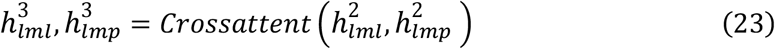

### Training configuration

For GraphKcat, we simultaneously predict the logKcat and logKm. The training or test entries either without logKcat or logKm values were masked. For DLkcat, and UniKP, we trained two models for logKcat and LogKm prediction. We used the code from https://github.com/SysBioChalmers/DLKcat for training DLkcat, code from https://github.com/Luo-SynBioLab/UniKP for training UniKP using the same training set of GraphKcat with their default parameters. All the training parameters can be found in **Supplementary Table S4**.

## Data availability

The PDBbind data set and CASF-2016 benchmark are available at http://www.pdbbind.org.cn. The processed data for training GraphKcat are available at https://doi.org/10.5281/zenodo.15396534. The TAL are collected from https://doi.org/10.1093/bib/bbae387.The DpADA are collected from https://doi.org/10.1021/acscatal.

## Code availability

The code and execution details for GraphKcat are available at https://github.com/DingLuoXMU/GraphKcat.

## Acknowledgments

This work has been supported by Scientific Research Innovation Capability Support Project for Young Faculty (ZYGXONJSKYCXNLZCXM-B6), National Natural Science Foundation of China (22121001), and Fundamental Research Funds for the Central Universities (20720240124).

