## Supplementary material for "Catalytic pocket-informed augmentation of enzyme kinetic parameters prediction via hierarchical graph learning": SI files

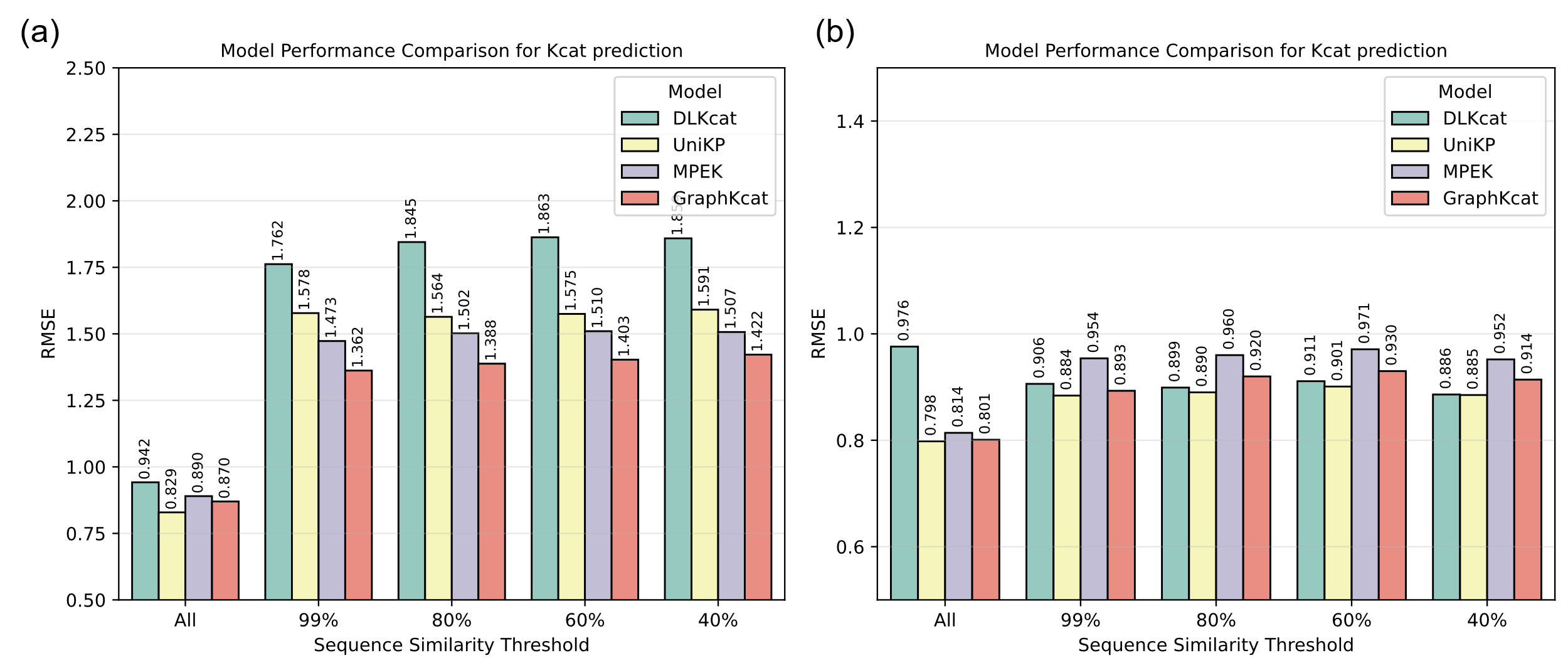

**Supplementary Figure 1**: **Comparative model performance**. (a-b) Root Mean Square Error (RMSE) for Kcat (a) and Km (b) predictions across DLKcat, UniKP, MPEK, and GraphKcat models under declining enzyme sequence similarity thresholds (100-40%) relative to the training set. All models were trained and evaluated on identical dataset.

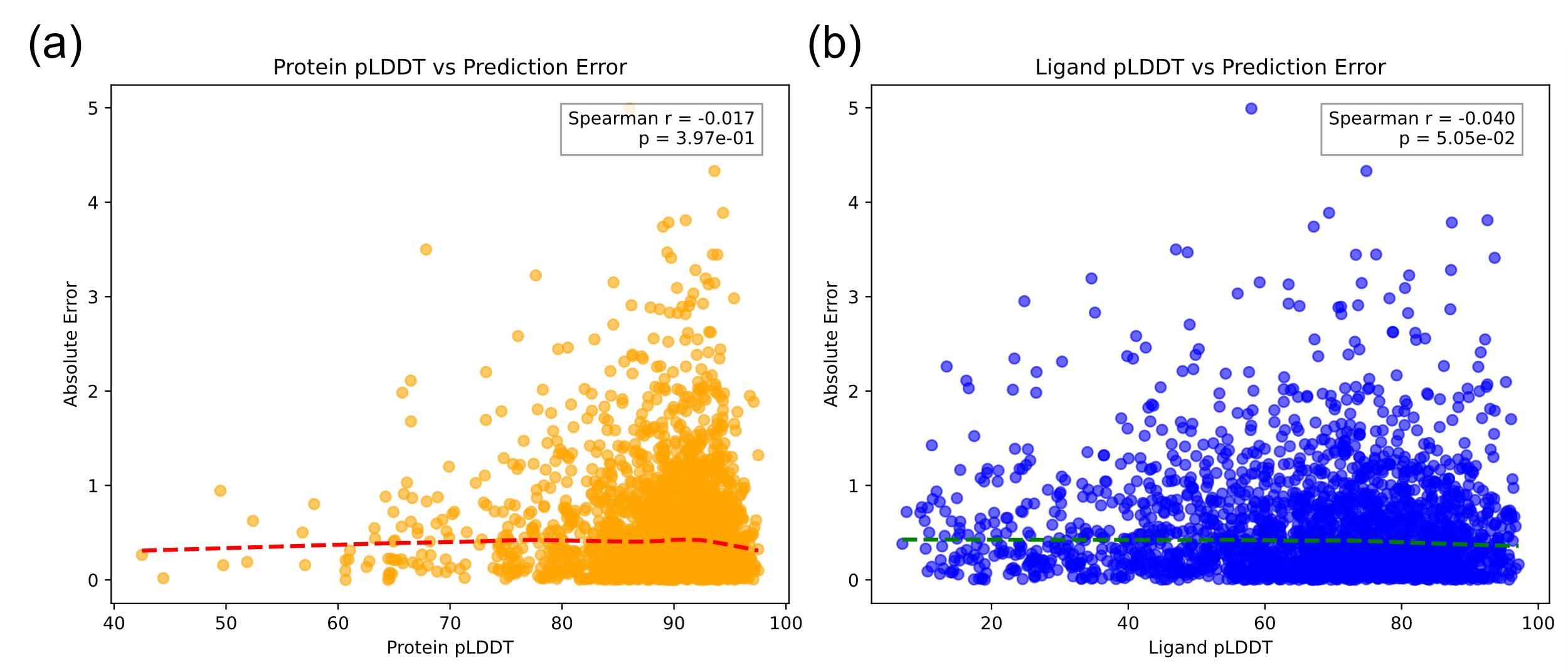

**Supplementary Figure 2**: **Spearman correlations between protein and ligand pLDDT values and absolute prediction errors for Km values prediction.** (a) for Protein and (b) for ligand.

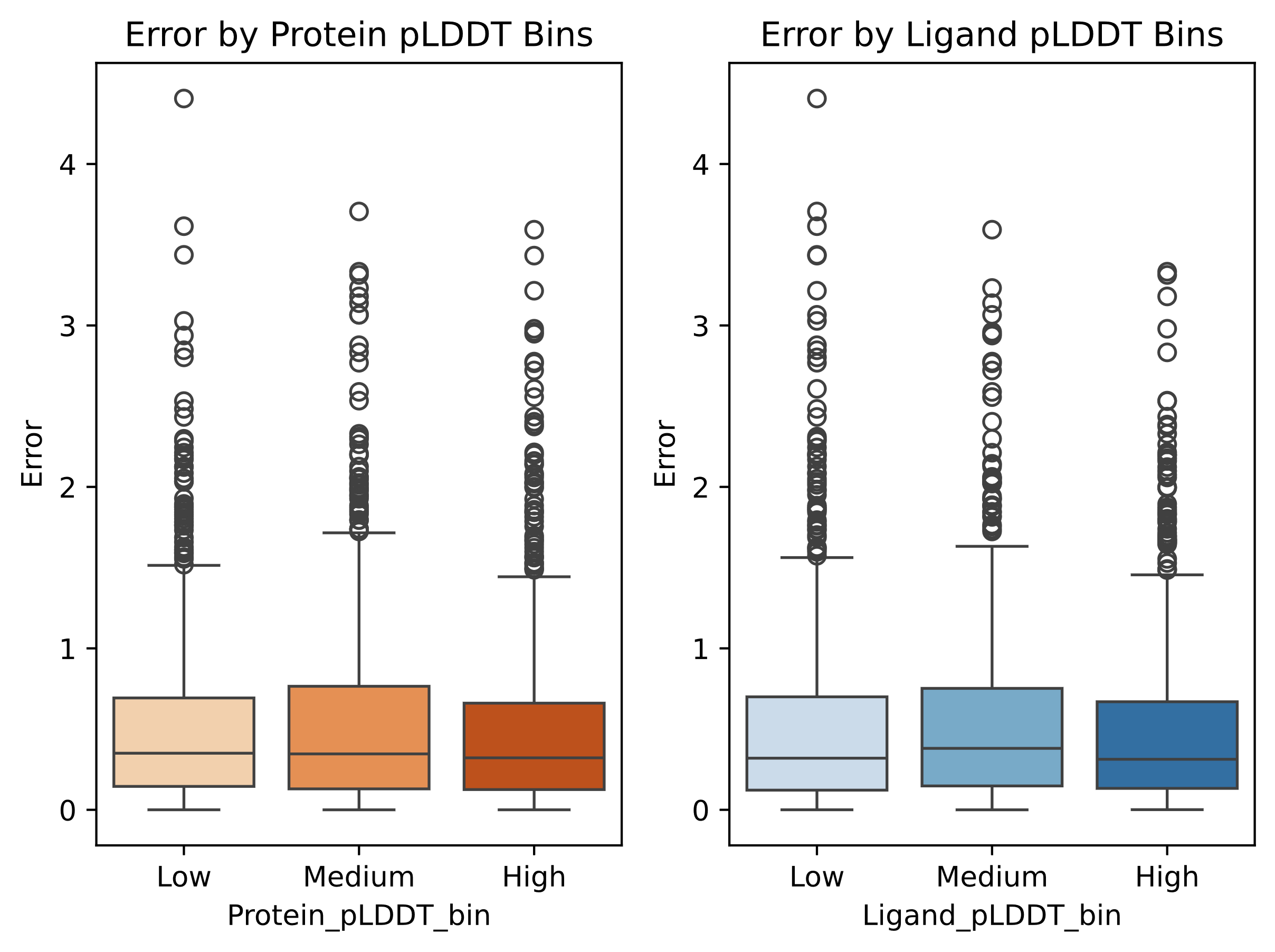

**Supplementary Figure 3**: **Prediction error distributions stratified by low/medium/high protein-ligand pLDDT confidence tiers for Km values prediction.** (a) for Protein and (b) for ligand.

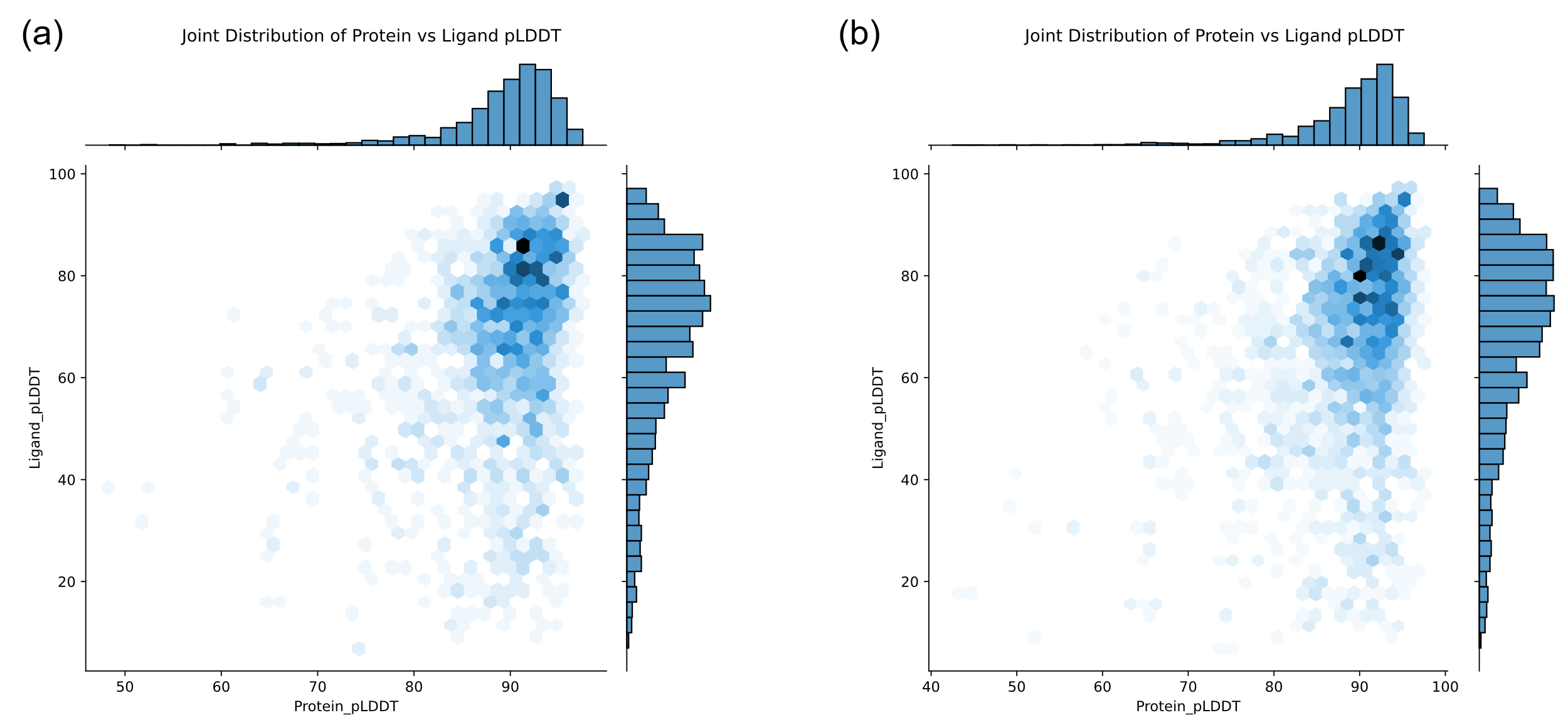

**Supplementary Figure 4**: **Joint protein-ligand pLDDT distribution for the predicted structures in Kcat entries (a) and Km entries (b) in test set.**

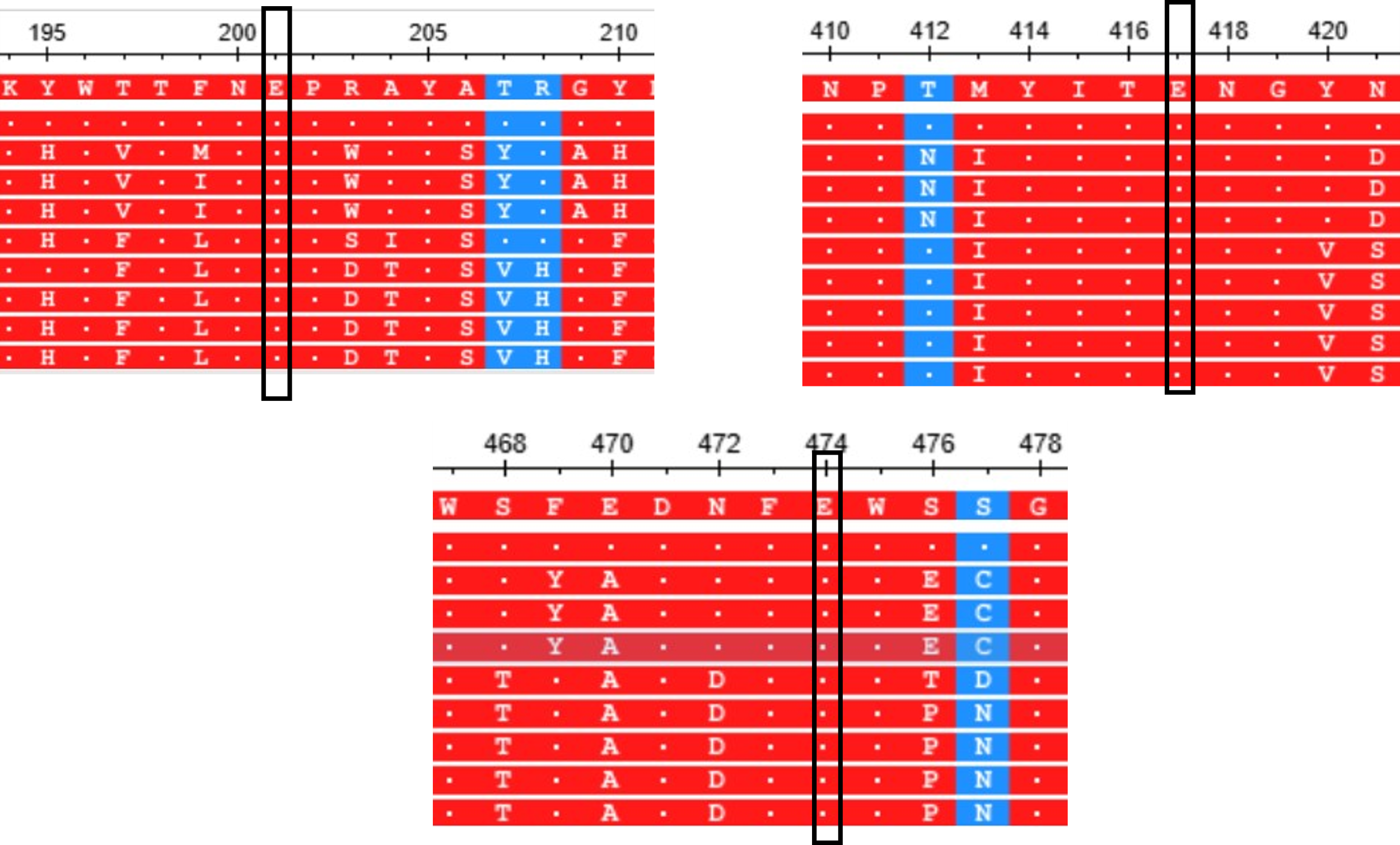

**Supplementary Figure 5**: multi-sequence alignment of β-galactosidase. Three conversed GLU (E201, E417, E474) were located in the enzyme active site. The MSA analysis were performed using the MSA viewer in Blast webserver.^1^

**Supplementary Table S1**: Comparison of the performance of machine learning scoring functions in CASF-2016 core set.

| Training set | Model | Algorithm | Size | Rp | RMSE |
| --- | --- | --- | --- | --- | --- |
| PDBbind general  set v.2016 | Graph Encoder | GNN | 11693 | **0.844** | **1.158** |
|  | GIGN^2^ | GNN | 11904 | 0.840 | 1.190 |
|  | IGN^3^ | GNN | 8298 | 0.837 | 1.220 |
|  | EGNN^4^ | GNN | 11904 | 0.816 | 1.289 |
|  | MPNN^2^ | GNN | 9662 | 0.813 | 1.511 |
|  | OnionNet^5^ | CNN | 11906 | 0.816 | 1.278 |
|  | Pafnucy^6^ | CNN | 11906 | 0.780 | 1.420 |

**Supplementary Table S2**: pLDDT values of predicted structures in TAL and DpADA enzyme mining dataset, and TAL enzyme mutation dataset. All those predicted structures were relaxed by Rosetta FastRelax module.

| Dataset | Entry | Protein pLDDT | Ligand pLDDT |
| --- | --- | --- | --- |
| TAL mining | AaTAL | 14.15 | 80.15 |
|  | RgTAL | 59.05 | 87.75 |
|  | IsTAL | 15.89 | 82.83 |
|  | SsTAL | 52.76 | 86.55 |
|  | HiTAL | 51.99 | 71.35 |
|  | LeTAL | 54.23 | 72.06 |
|  | PcTAL | 82.68 | 76.94 |
| TAL mutation | MT-10Y | 60.09 | 87.38 |
|  | MT-603P | 60.30 | 87.82 |
|  | MT-366W | 56.55 | 87.96 |
|  | MT-337C | 60.83 | 87.86 |
|  | MT-668S | 61.83 | 87.93 |
|  | MT-489T | 62.93 | 87.35 |
|  | MT-337D | 58.32 | 87.61 |
|  | MT-587V | 59.95 | 88.04 |
|  | MT-366H | 42.82 | 87.45 |
| ADA mining | DpADA | 31.46 | 88.88 |
|  | ADA1 | 23.02 | 84.82 |
|  | ADA2 | 22.19 | 86.69 |
|  | ADA3 | 18.93 | 87.42 |
|  | ADA4 | 21.35 | 87.25 |
|  | ADA5 | 23.01 | 87.79 |
|  | ADA6 | 24.45 | 87.39 |
|  | ADA7 | 27.05 | 90.27 |
|  | ADA8 | 21.39 | 87.78 |
|  | ADA9 | 26.76 | 90.65 |
|  | ADA10 | 25.66 | 90.68 |

**Supplementary Table S3**: node and edge feature description of GNN models

| All-atom graph | **Features** | Dim^a^ | Description |
| --- | --- | --- | --- |
|  | **Nodes** |  |  |
|  | Atom type | 10^b^ | Heavy atom type [C, N, O, S, F, P, Cl, Br, I, others] |
|  | Degree | 7^b^ | Number of covalent bonds [0,1,2,3,4,5,6,] |
|  | Implicit valence | 7^b^ | Implicit valence of the atom [0,1,2,3,4,5,6] |
|  | Hybridization | 5^b^ | [sp, sp2, sp3, sp3d, sp3d2] |
|  | Aromatic | 1 | Whether the atom part of an aromatic system |
|  | Hydrogens | 5 | Number of connected hydrogens [0,1,2,3,4] |
|  | **Edges** |  |  |
|  | Edge type | 3^b^ | Edge type of ligand atom-ligand atom, ligand atom-protein atom, protein atom-protein atom [0,1,2] |
|  | Distance | 1 | Distance of atom-atom |
|  | Coordinate | 3 | x, y, z |
| Coarse-grained graph | **Nodes(ligand-ligand)** |  |  |
|  | Subgraph type | 350^b^ | One hot encoding of ligand subgraph types (1427+ Unknown) |
|  | Pooled features | 256 | Pooled features of ligand nodes form all-atom graph |
|  | **Edges(ligand-ligand)** |  |  |
|  | Subgraph distance | 16* | The encoding of the centroids - centroids distance of subgraph nodes  in terms of Gaussian radial basis functions |
|  | **Nodes(protein-protein)** |  |  |
|  | Sequence | 20 | A one-hot representation of amino acid identity |
|  | Dihedrals | 6 | {sin, cos} ◦ {φ, ψ, ω} computed from Ci−1, Ni, Cαi, Ci, and Ni+1 |
|  | Pooled features | 256 | Pooled features of protein nodes form all-atom graph |
|  | **Edges(protein-protein)** |  |  |
|  | Amino acid distance | 16 | The encoding of the C*α_j_* − C*α_i_* distance between amino acids nodes  in terms of Gaussian radial basis functions |
|  | Relative position | 16 | Relative position-based sin/cos position encoding of each edge |
|  | **Edges(ligand-protein)** | 16 | The encoding of the centroids- C*α* distance between ligand subgraph and amino acids nodes  in terms of Gaussian radial basis functions |

^a^: Dimension of feature vector.

^b^: One hot encoding of features.

**Supplementary Table S4**: The hyperparameter setting of each model.

| Model | Hyperparameters | Setting |
| --- | --- | --- |
| EGNN model | Number of layers | 4 |
|  | Attention head | 4 |
|  | Hidden dimension | 256 |
|  | MLP layers | 2 |
|  | Dropout | 0.2 |
|  | Learning rate | 1e-4 |
|  | Weight decay | 1e-6 |
|  | Batch size | 8 |
|  | Epochs | 1000 |
|  | Optimizer type | AdamW |
|  | Pooling | mean |
|  | Vocab size | 350 |
|  | Share MLP parameters | False |
